# Multimodal interferometric imaging of nanoscale structure and macromolecular motion uncovers UV induced cellular paroxysm

**DOI:** 10.1101/428383

**Authors:** Scott Gladstein, Luay M. Almassalha, Lusik Cherkezyan, John E. Chandler, Adam Eshein, Aya Eid, Di Zhang, Wenli Wu, Greta M. Bauer, Andrew D. Stephens, Simona Morochnik, Hariharan Subramanian, John F. Marko, Guillermo A. Ameer, Igal Szleifer, Vadim Backman

## Abstract

We present a multimodal label-free interferometric imaging platform for measuring intracellular nanoscale structure and macromolecular dynamics in living cells with a sensitivity to macromolecules as small as 20nm and millisecond temporal resolution. We validate this system by pairing experimental measurements of nanosphere phantoms with a novel interferometric theory. Applying this system *in vitro*, we explore changes in higher-order chromatin structure and dynamics that occur due to cellular fixation, stem cell differentiation, and ultraviolet (UV) light irradiation. Finally, we discover a new phenomenon, cellular paroxysm, a near-instantaneous, synchronous burst of motion that occurs early in the process of UV induced cell death. Given this platform’s ability to obtain nanoscale sensitive, millisecond resolved information within live cells without concerns of photobleaching, it has the potential to answer a broad range of critical biological questions about macromolecular behavior in live cells, particularly about the relationship between cellular structure and function.

## Introduction

At the level of individual living cells, thousands of unique molecules are constantly moving, interacting, and assembling-working to execute cellular functions and keep the cell alive. Understanding the properties of this complex motion and its interplay with the cellular ultrastructure remains one of the most critical and challenging topics of study in modern biology. While widely explored, the link between nanoscale structure and molecular motion is particularly challenging to study for several reasons: (1) nanoscale macromolecular organization is often composed of hundreds to thousands of distinct molecules, some of which cannot be easily labeled such as lipids, nucleic acids, or carbohydrates, (2) molecular dynamics depends uniquely on the timescales of interest in the context of the surrounding macromolecular nanostructure, and (3) molecular motion and ultrastructure evolve in concert but along distinct timescales, often spanning milliseconds to days.

Most techniques to study molecular motion in eukaryotic cells require the use of exogenous small molecule dyes or transfection-based fluorophore labeling. These techniques, such as single molecule tracking, fluorescence recovery after photobleaching (FRAP)^1,2^, photoactivation^3,4^, fluorescence correlation spectroscopy (FCS)^5^, and F□rster resonance energy transfer (FRET)^6^ have greatly expanded our understanding of the behavior of molecular motion in live cells. Despite their utility and the insights produced regarding cellular behavior, these methods have limitations. For instance, single molecule tracking, FRET, and FCS provide information on the activity of individual molecules, but cannot probe the motion of complex macromolecular structure that often govern cellular reactions, such as the supra-nucleosomal remodeling that may occur during gene transcription or DNA replication. Likewise, FRAP and photoactivation can yield diffraction-limited information about the general molecular mobility within cellular compartments, but requires the use of high intensity photobleaching which may damage the underlying structure.

Beyond technique specific applications, these methods share common limitations: (1) they can only probe the behavior of an individual or a few molecules concurrently; (2) they require the use of either potentially cytotoxic small molecule dyes or transfection, which often cannot label lipid or carbohydrate assemblies directly; (3) they are susceptible to artifacts due to photobleaching; and (4) they have significant limitations to probe cellular heterogeneity due to the inherent variability of label penetrance, a critical feature of multicellular systems and diseases, including cancer^7-9^. Further, to extend these techniques to study the interplay between local structure and motion requires the use of additional fluorophores, which have similar drawbacks.

To address these issues, we present a novel label-free interferometric platform that captures the temporal behavior of macromolecular assemblies in live cells by building upon live cell Partial Wave Spectroscopy (PWS), a quantitative imaging technology that provides label-free measurements of nanoscale structure^10^. PWS obtains this information by taking advantage of an interference phenomenon in the light backscattered from intracellular macromolecular structures. This interference produces spectral variations that depend on the nanoscale organization of these structures. PWS has resulted in many breakthroughs in the study of the higher-order organization of chromatin structure, its relation to the development of cancer^7,11^, and its use in cancer diagnostics^12–17^ and therapeutics^18^. The same interference phenomenon that enables PWS to probe intracellular structure at length-scales below the diffraction limit without the use of labels can be utilized to retrieve information on macromolecular motion by measuring temporal variations in the interference signal instead of spatial (wavelength) variations. By combining these two techniques, we pair measurements of cellular dynamics with macromolecular structure – creating a dual light interference platform (dual-PWS) to greatly enhance our ability to probe cellular behavior at the level of macromolecular assemblies. While dual-PWS is not molecularly specific, it captures the underlying behavior of all macromolecular assemblies and is complementary to the aforementioned molecular specific techniques^10^. Additionally, as this method relies on analysis of scattered light, its temporal limitations are defined by current optical technology (illumination intensity, detector sensitivity, etc.) and not by the photochemical limits of probe dyes. Consequently, this interferometric platform measures nanoscale macromolecular motion in tandem with spatio-temporal behavior of the macromolecular ultrastructure in dozens of live cells simultaneously without photobleaching artifacts.

Herein, we present the theory for temporal interference and validate this system using experimental measurements of nanosphere phantoms. Applying dual-PWS microscopy *in vitro*, we explore nanoscale structural and dynamic changes that occur due to cross-linking chemical fixation and the differentiation of stem cells. Then, we probe the spatio-temporal response of macromolecular assemblies to ultraviolet (UV) light. We discover a new phenomenon that occurs early in the process of UV induced cell death: a near-instantaneous burst of motion, referred to in this manuscript as cellular paroxysm. These paroxysms are asynchronous between cells, are uncorrelated with membrane permeabilization, and are directly correlated with phosphotidylserine (PS) externalization and depolymerization of the actin cytoskeleton. This nanoscale transformation is synchronous across the cell, originating at loci that are microns apart within 35 milliseconds. This cellular paroxysm indicates the existence of an undiscovered phenomenon that may play a role in the earliest stages of cell death. Altogether, we demonstrate that dual-PWS measures nanoscale structure and dynamics with millisecond temporal capabilities, allowing for the label-free quantification of macromolecular motion and structure in live cells. These new capabilities enable the discovery of new phenomena not previously possible by other techniques.

## Results

### Theory: Origin of Back-Scattered Interference

The dual-PWS system takes advantage of light interference to amplify a weak scattering signal from cellular structures, allowing us to probe the nanoscale structural organization and dynamics without necessitating exogenous labels in live cells^10,19^. The interference signal originates from (1) a strong reflectance from the glass-cell interface (petri dish) due to the high refractive index (RI) mismatch, referred to as the reference arm, and (2) scattering from within the cell due to refractive index fluctuations, referred to as internal scattering. This interference signal will vary spectrally, referred to in this paper as spectral interference, depending on the organization of intracellular structures, which generate the internal scattering. Similarly, this interference signal will vary in time, referred to in this paper as temporal interference, depending on the motion of those intracellular structures.

### Theory: Analysis of Spectral Interference

PWS microscopy utilizes the spectral interference to measure the structural organization within cells. PWS quantifies the variations in mass density within a sample, Σ*_s_*, (i.e. the heterogeneity of the macromolecular organization) with a sensitivity to length-scales ranging from 20–200nm. Σ*_s_* is acquired by calculating the standard deviation of the spectral interference. Simulations, further biological explanations, and details on measurement and theory of Σ*_s_* can be found in previous works^10,19–23^.

### Theory: Analysis of Temporal Interference

Several properties of macromolecular motion are simultaneously obtained by capturing the backscattered temporal interference spectrum, of which the two most prominent are (1) the fractional moving mass and (2) the correlation time/diffusion coefficient.

The fractional moving mass, *m_f_*, is the product of the mass of the typical moving macromolecular cluster, *m_c_*, and the volume fraction of mobile mass, Φ, within the sample. This property is acquired by measuring and normalizing the variance of the temporal interference, 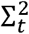.

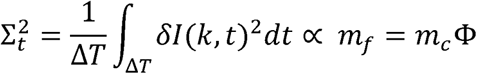

where δI(k,t) is the time-varying part of the intensity reflectance at wavenumber, k, and ΔT is the acquisition time. In order to better understand these parameters, examples of the effect of different *m_c_* and Φ on fractional moving mass (*m_f_*) are explored. In a healthy cell, chromatin moves in a highly dynamic manner, continuously altering its structure and function (large Φ, large *m_f_*). However, the addition of chemically crosslinking paraformaldehyde would decrease the fractional moving mass by decreasing the volume fraction of moving chromatin without changing the density or length-scales (same *m_c_*, small Φ, small *m_f_*). Next, a comparison of the effect of molecular length-scales on the fractional moving mass can be made by considering the motion of actin filament bundles in comparison to an equivalent amount of actin monomers. Even though the volume fraction and total mass is equivalent in both samples, each moving structure in the filament sample is larger, and therefore produces a greater fractional moving mass (large *m_c_*, large *m_f_*) compared to the sample of monomers (small *m_c_*, same Φ, small *m_f_*). It should be noted that dual-PWS does not specifically measure actin filaments, individual chromatin loops, etc.; instead, it measures the physical and dynamic properties of all macromolecular assemblies at each location inside the cell within the sensitivity range.

In contrast to the fractional moving mass, *m_f_* (quantified through the variance of the temporal interference), the autocorrelation of temporal interference describes how correlated the motion is at different timescales. Physical properties, such as the diffusion coefficient, *D*, can be extracted from the shape of the temporal autocorrelation function. While not all motion within the cell is specifically diffusive, *D* provides a convenient parameter for measuring the rate of the macromolecular motion. The diffusion coefficient, *D*, is acquired by calculating the decay rate of the temporal autocorrelation function as follows

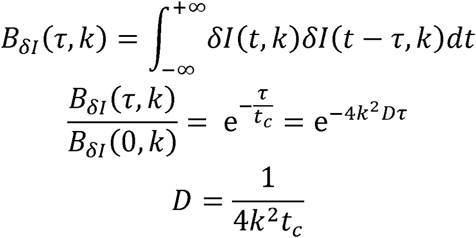

where B_δI_(k,t) is the autocorrelation of δI, τ is the time lag of the autocorrelation function, and t_c_ is the correlation time. A more detailed derivation of the theory behind *m_f_* and *D* can be found in the **SI Temporal Interference Theory**.

The system’s sensitivity range to molecular motion will depend on the exposure time of each frame for the fastest processes and the length of acquisition for the slowest processes. For a 32ms exposure and 6.4s acquisition time (parameters used in phantom experiments), our system is sensitive to processes with diffusion coefficient, 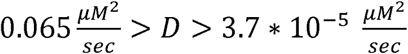 (*see* **SI Timescale Sensitivity**). This results in a sensitivity to a broad array of biological processes within the nucleus, including the diffusion of genomic loci 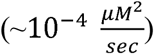^24^, transcription factors bound to DNA 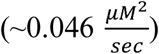^25^, and mRNA 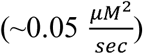^26^. Theoretically, the fractional moving mass for isochronic motion is independent of the speed of the moving material, while in practice the measurements will be most sensitive to processes in the middle of the sensitivity range. The system will detect all non-isochronic motion regardless of the speed, but they may not be fully quantifiable as the full signal is not resolved. It should be noted that there are no physical limitations to the speed of processes to which dual-PWS is sensitive. The lower bound of the sensitivity range can be easily adjusted by acquiring more or fewer frames in each acquisition. The sensitivity to faster processes can be improved with currently available hardware modifications to increase the SNR and lower the exposure time (*see* **SI High Temporal Resolution System**).

### Nanosphere Phantom Validation

To validate the nanoscale sensitivity of the dual-PWS system to molecular motion, a phantom was constructed by suspending carboxyl functionalized nanospheres (Phosphorex Inc.) in a glycerol-water solution (90% glycerol RI = 1.46 or 70% glycerol RI=1.44). The glass bottoms of the petri dishes were replaced with sapphire (RI=1.76) to maintain a similar refractive index mismatch for our reference arm (glass-media vs. sapphire-glycerol) in comparison to live cell experiments. Spheres of different sizes and volume fractions were tested to validate physical properties that underlie changes in the fractional moving mass and diffusion in cells (**Figure 1**). For fractional moving mass, spheres with radii of 100nm, 50nm, 37.5nm, and 25nm were used to vary *m_c_*, the mass of the typical moving macromolecular cluster, while sphere solutions of different volume fractions (0.1% spheres in 90% glycerol vs. 0.3% spheres in 70% glycerol) were compared to represent changes in Φ. For the most accurate system validation, experimental measurements are compared to the analytical expression of Σ*_t_* (from eq. 6, where τ = 0), which removes some of the assumptions used in deriving *m_f_*. Additional corrections to account for exposure time are applied (*see* **SI Timescale Sensitivity**). As expected Σ*_t_ is sensitive to changes* in both *m_c_* and Φ, closely matching theory [∆Σ*_t_* = 21.5 ± 3.5% (SEM), n = 5, averaged percent error between experiment and theory, R^2^ = 0.78] (**Figure 1b**). For validation of diffusion coefficients, *D*, experimental measurements of the 0.1% spheres at different sizes are compared to theoretically predicted values using the Stokes-Einstein equation; the measured and theoretical values closely matched for 0.1% solutions [*ΔD* = 16.9 ± 4.1% (SEM), n = 5, averaged percent error between experiment and theory, and R^2^ = 0.993] (**Figure 1c**). As expected, diffusion measurements of the 0.3% solutions are not accurate as the decreased viscosity and exposure time limits result in sphere speeds outside our sensitivity range (**SI Figure 3**). This loss in sensitivity is compensated for in the Σ*_t_* comparison using exposure time correction, but the same cannot be done for measurements of diffusion. Although these spheres are not visible using a traditional wide-field microscope due to the diffraction limit, the interference with our reference amplifies the scattering signal, allowing visualization of these nanospheres and measurement of their dynamic properties using dual-PWS (**SI Figure 6**).

**Figure 1:**
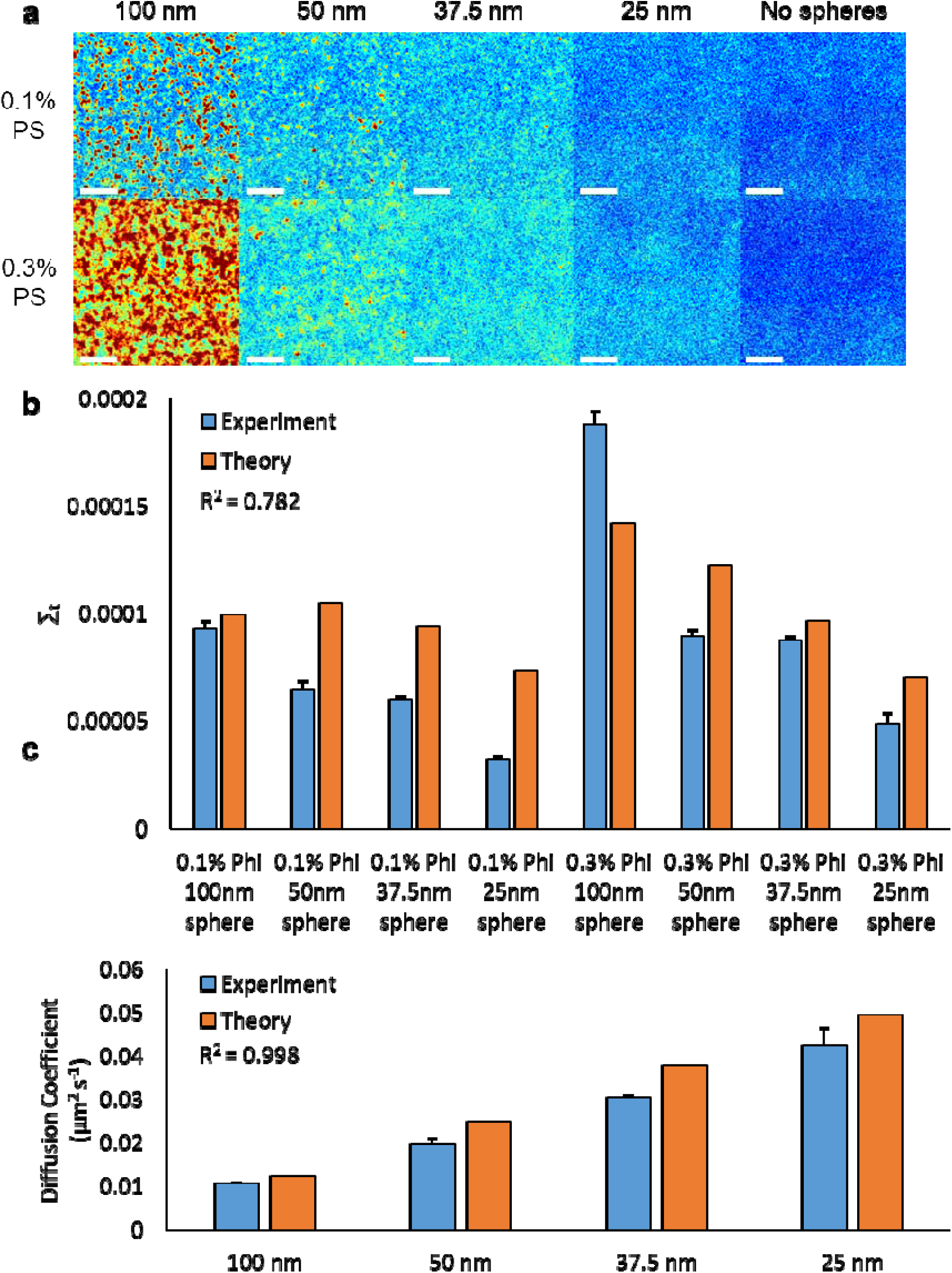
Nanosphere Phantoms. **a)** Representative ∑_t_ maps of polystyrene nanosphere phantoms created with a variety of nanosphere sizes (25nm, 37.5nm, 50nm, and 100nm) and concentrations (0.1% and 0.3% volume fractions). **b)** Bar graph comparing ∑_t_ ± SEM for the phantoms represented in (a) with analytical theory. The PWS measurements of these phantoms match well with theory (R^2^=0.782) and show that ∑_t_ is sensitive to the volume fraction of spheres (φ) and the mass of the moving spheres (m_c_). **c)** Bar graph comparing experimentally measured *D* ± SEM with theoretical values calculated using the Stokes-Einstein equation for 0.1% sphere phantoms with sizes 25nm, 37.5nm, 50nm, and 100nm. The PWS measured *D* values closely match the predicted values for a correlation of R^2^ = 0.998. Scale bar is 8 uM.

### Cellular Fixation

To confirm the label-free capacity to obtain information on both cellular dynamics and structure in live cells, we measured the effects of cellular fixation in HeLa cells (**Figure 2a**). Under normal conditions, cells demonstrate considerable macromolecular motion throughout the nucleus and the cytoplasm. While living cells exhibit a large fractional moving mass, motion disappears upon cross-linking chemical fixation [10 minutes, 3.6% paraformaldehyde solution; nucleus, Δ*m_f_* = −97.9 ± 0.8% (SEM), n = 7, p-value = 0.0006, paired comparisons; cytoplasm, Δ*m_f_* = −95.3 ± 1.7% (SEM), n = 7, p-value = 0.0018, paired comparisons]. Within this experiment, chemical fixation induced alterations in cytoplasm structure [∆Σ*_s_* = 33.5 ± 2.7% (SEM), n = 7, p-value = 0.0003, paired comparisons], but preserved the average structural heterogeneity within the nucleaus [∆Σ*_s_* = −2.3 ± 3.5% (SEM), n = 7, p-value = 0.55, paired comparisons] (**Figure 2b**). A more thorough examination of the effects of fixation on nanoscale structural heterogeneity has been reported in previous publications^27,28^.

**Figure 2:**
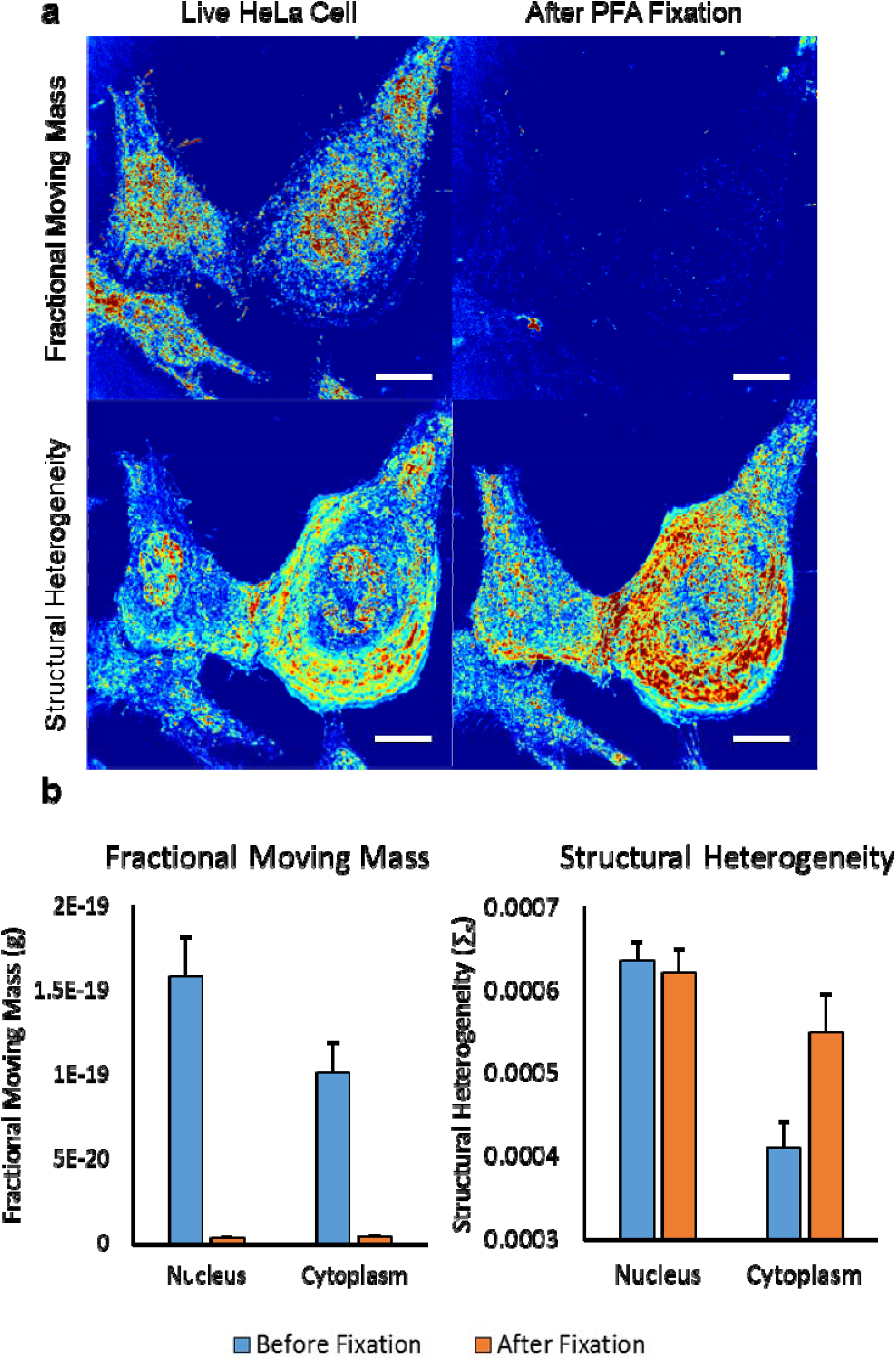
Cellular Fixation. **a)** Representative m_f_ (top) and ∑_s_ (bottom) maps of HeLa cells before and after crosslinking fixation using paraformaldehyde (PFA). **b)** Bar graph quantifying the mean m_f_ ± SEM and ∑_s_ ± SEM for the nucleus and cytoplasm separately. Scale bar is 20 uM.

### Stem Cell Differentiation

Human mesenchymal stem cells (hMSCs) differentiate into different specialized cell types, including osteoblasts, due to the integration of a host of chemical and physical signals. The process of cellular differentiation is among the most widely studied in molecular biology and has been demonstrated to result in changes in the organization of chromatin in fixed cells as well as the motility of transcription factors^29,30^. While alterations in the dynamics of individual transcription factors have been observed, no one has studied large scale changes in motion due to differentiation. We hypothesized that in hMSCs there would be increased chromatin heterogeneity and higher mobility due to the unique abilities of hMSCs to access genes from multiple phenotypes and have increased activity due to the process of differentiation. Consistent with previous studies, we observe that the hMSC and osteoblast populations have very different states of chromatin folding and nuclear dynamics (**Figure 3a**) as well as significant heterogeneity within each population (**SI Figure 7**). Specifically, hMSCs were observed to have a more heterogeneous chromatin structure [∆Σ*_s_* = 25.0 ± 2.2% (SEM), p-value = 9.6e-24], a higher fractional moving mass [Δ*m_f_* = 24.8 ± 3.9% (SEM), p-value = 8.8e-10)], and a faster molecular motion [Δ*D* = 6.1 ± 2.6% (SEM), p-value = 0.017] when compared to differentiated osteoblasts (n=166 hMSC and n=102 osteoblasts) (**Figure 3b**).

**Figure 3:**
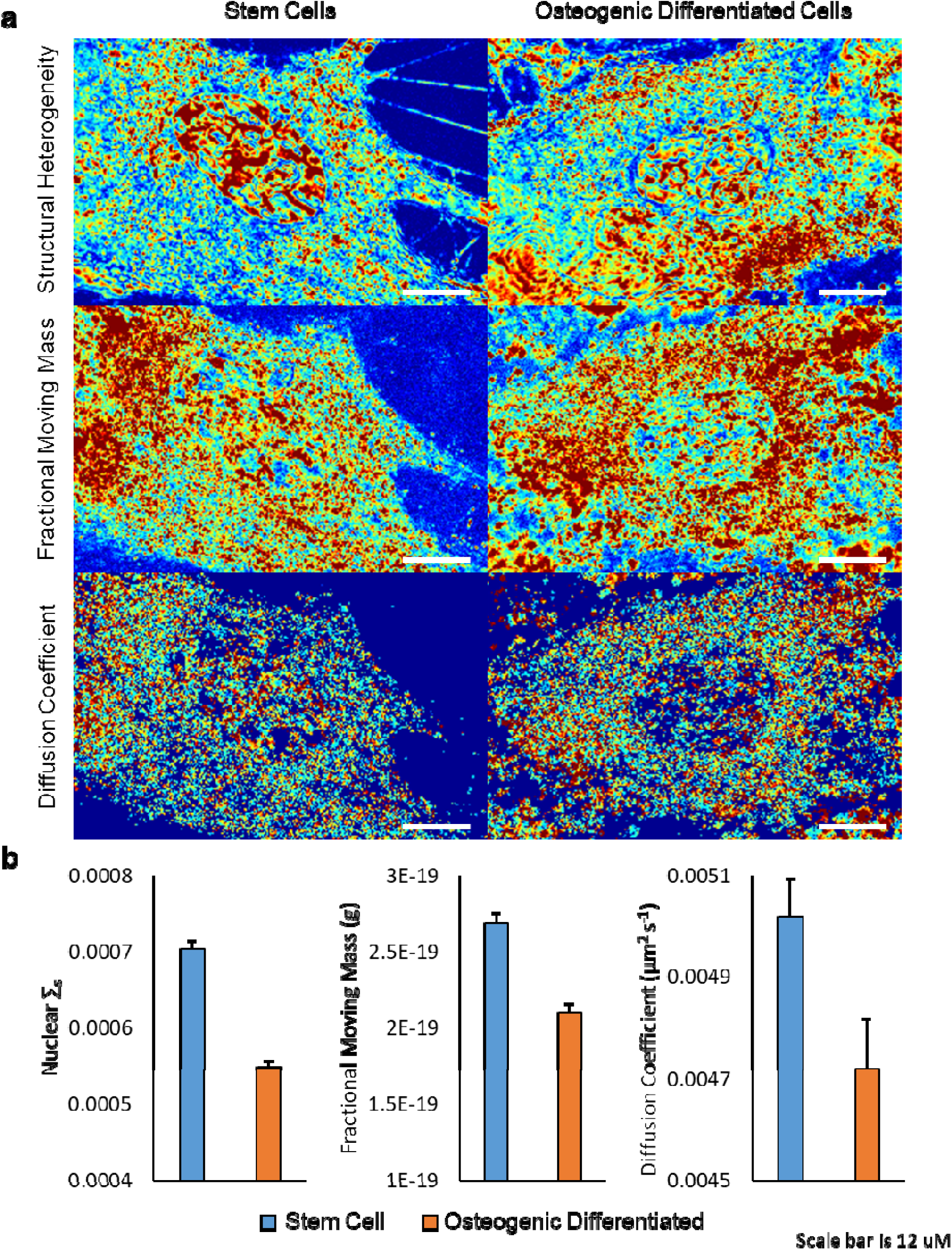
Stem Cell Differentiation. **a)** Representative ∑_s_ (top), m_f_ (middle), and *D* (bottom) maps of human mesenchymal stem cells (left) and osteoblasts (right). **b)** Bar graph quantifying the mean ∑_s_ ± SEM, m_f_ ± SEM, and *D* ± SEM for the stem cells and osteoblasts. We observe different states of chromatin folding and nuclear dynamics comparing hMSCs to osteoblasts. Scale bar is 12 uM.

### Ultraviolet Irradiation

While UV light is most frequently associated with DNA damage^31^, it also has wide ranging effects on the cellular membrane^32^, mitochondrial behavior^33^, and chromatin structure^10^. To date, not much is known about the real-time effects on nanoscale (20–350nm) cellular structure and dynamics due to limitations in existing microscopic techniques as UV illumination can distort the optical properties of dyes as well as induce significant structural damage to cells. Previous work with live cell PWS microscopy to detect alterations in higher-order chromatin structure^34^ after UV irradiation demonstrated previously unobserved transformation in cellular and chromatin ultrastructure in real-time. In this study, we apply our dual-mode system to monitor both the transformation to cellular ultrastructure and macromolecular dynamics during UV illumination. To perform UV irradiation, the cells were illuminated with the full lamp spectra, resulting in UV irradiation of ~0.013 μW/μm^2^ between 350 and 400nm. For control measurements, a UV filter is placed in the system to block all light below 405nm. In control cells, the structure and diffusion coefficients are stable throughout the 25 minute experiment; the fractional moving mass is mostly stable, but does show a slight decrease [Nucleus: Δ*m_f_* = −12.9 ± 7.0% (SEM), n = 28, p-value = 0.003; Cytoplasm: Δ*m_f_* = −6.4 ± 2.7% (SEM), n = 28, p-value = 0.001; paired comparisons between time point t = 0 and t = 25 min] (**Figure 4ab**). While the structure on average remains stable, localized structure continuously evolves consistent with the large fractional moving mass detected with the temporal measurements (**Video 1**). At the level of the nucleus, this indicates that even at very short timescales (milliseconds), supra-nucleosomal organization is continuously evolving.

**Figure 4:**
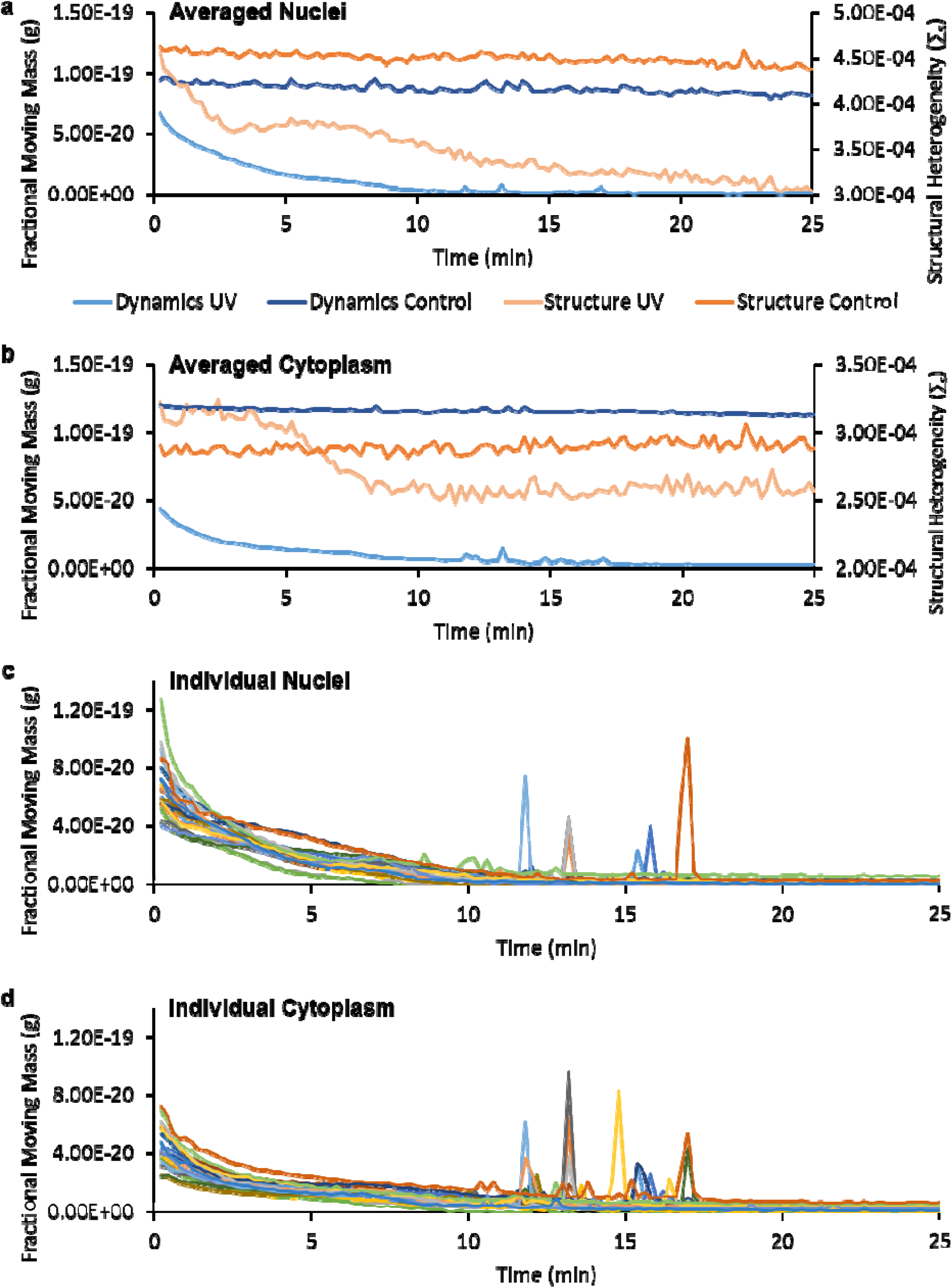
Ultraviolet Irradiation Quantification. Line graphs showing the averaged ∑_s_ and m_f_ in response to UV (n=20) and non-UV (n=28) irradiation within the nucleus **(a)** and the cytoplasm **(b)**. Line graphs showing individual cell response to UV irradiation within the nucleus **(c)** and the cytoplasm **(d)**. By examining individual cells, a near-instantaneous, stochastic burst of motion, termed a cellular paroxysm, occurs between 10 and 20 minutes that is asynchronous from cell to cell.

Interestingly, continuous UV irradiation induces a steady decrease in the fractional moving mass (↓*m_f_*) within the first ten minutes of irradiation, ultimately stabilizing in a state of low dynamics for the following fifteen minutes. Structurally, this attenuation in the fractional moving mass is first accompanied by a sharp homogenization in chromatin organization (↓Σ*_s_*) for the first three minutes, followed by a variable duration plateau spanning from three to eight minutes that precedes a further homogenization of chromatin structure (↓Σ*_s_*) for the remainder of the experiment (**Figure 4a**). The diffusion coefficient shows a decrease in the rate of molecular motion after the first four minutes of UV irradiation; the diffusion coefficient was not analyzed for the entirety of the experiment due to low SNR at later time points (*see* **SI UV Diffusion Analysis; SI Figure 5**). The structural response in the cytoplasm is distinct from the nuclear response. Within the cytoplasm, there are no structural alterations for the first ~4.5 minutes. Then, there is a rapid homogenization of structure (↓Σ*_s_*) for the next ~4.5 minutes, before the signal stabilizes for the remainder of the experiment (**Figure 4b**). Notably, once the cells reach their low dynamic state, the localized structural alterations stop and the spatial distribution of chromatin organization is fixed for the rest of the experiment (**Video 1**).

Taking advantage of capabilities of live cell PWS microscopy to analyze structure and dynamics in the same cell without photobleaching, we examine the temporal responses of individual cells due to UV irradiation (**Figure 4cd**). In particular, a near-instantaneous, stochastic burst of motion, termed a cellular paroxysm, occurs between 10 and 20 minutes that is asynchronous from cell to cell. Critically, this analysis is possible due to the dual-PWS system’s capacity to perform single cell tracking as we observe a unique stochastic phenomenon that is not apparent in the aggregate behavior of the population. From these measurements, it is clear that these bursts of activity generally occur within a single acquisition (< 7 seconds).

By examining the temporal interference (intensity as a function of time at each pixel) of each cellular event, we observe that the initiation of these bursts of motion can happen in as little as a single 35ms frame and can occur independently at distances over 30µm apart. To quantitatively analyze this phenomenon, we enhanced the temporal resolution to a single frame using spectral feature detection (**Figure 5a**) and localized the motion burst based on their temporal initiation (**Figure 5cd)**. **Figure 5b** shows a histogram of the timing of the event for each pixel within a single cell obtained using spectral feature detection (MATLB findchangepts function); surprisingly, we found that within this particular cell, following initiation, 1% of the events occur with the first 35ms, 19% within 70ms, and over 50% within 140ms. An interesting feature of this spatio-temporal organization is that these events do not originate in a single location and spread out in a wave throughout the cell as one might assume (**Figure 5cd** and **Video 2)**. The initial motion occurs simultaneously within 35ms on opposite ends of the cell and extends into the cell nucleus; to accomplish this, a molecular regulator synchronizing these events would need to diffuse at a rate 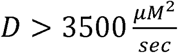. Consequently, this motion is orders of magnitude faster than protein diffusion 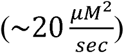 in the eukaryotic cytoplasm.

**Figure 5:**
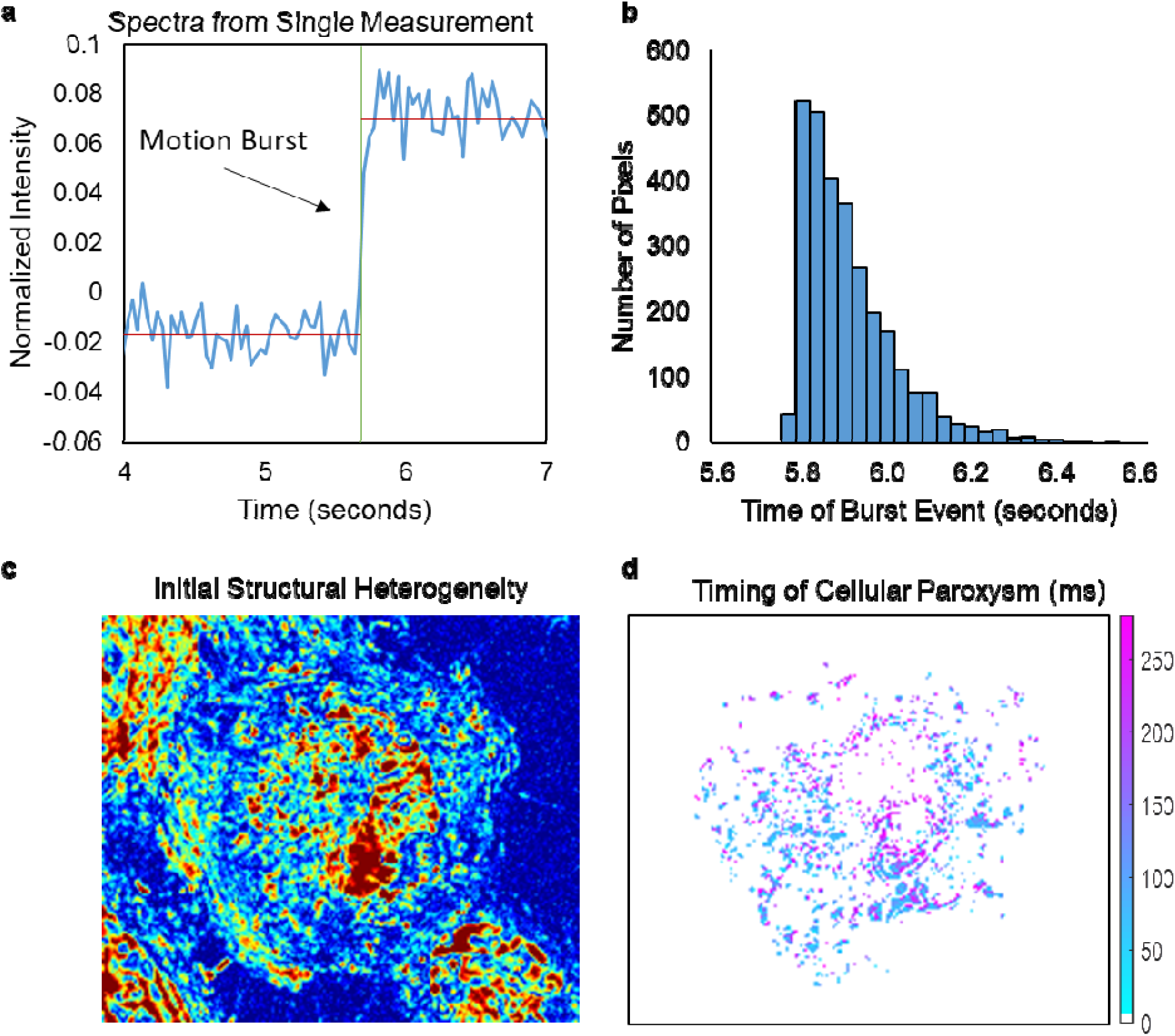
Cellular Paroxysm Temporal Analysis. **(a)** Example temporal spectra from a single pixel during a cellular paroxysm. Using spectral feature detection the timing of the event can be determined for each pixel reducing the temporal resolution from the full acquisition time (~7s) to a single exposure time (35ms). **(b)** Example histogram of the timing of the event for each pixel within a single cell. **(c)** ∑_s_ map of this cell at the beginning of the experiment. **(d)** Map displaying the timing of the event for each pixel within a representative cell. The spatial distribution of this timing shows that the initial motion occurs simultaneously within a single 35ms frame on opposite ends of the cell.

As this is a newly observed phenomenon, we explored molecular transformations that accompanied these cellular paroxysms. Owing to the stochastic nature of these events, UV irradiation of cells was performed for an intermediate duration in order to differentially induce the response in some, but not all, cells within the field of view. Using this approach, differential molecular analysis could be performed to examine the differences between cells that underwent the cellular paroxysm (referred to as CP cells) and cells that did not (referred to as NCP cells) under comparable UV illumination. As these bursts of activity occurred throughout the entire cell, we hypothesized that these events were occurring at the level of either the cytoskeleton (actin disruption) or the plasma membrane (pore formation or phosphatidylserine externalization). Alexa Fluor 488 Phalloidin, which binds to F-actin, enables imaging of the cytoskeletal structure. A decrease in Phalloidin fluorescence intensity was observed for cells irradiated with UV with a larger decrease in CP cells [-81.3 ± 0.7% (SEM); n = 18 (CP) and 22 (controls), p-value = 2.8e-17] compared to NCP cells [-71.0 ± 2.7% (SEM); n = 20 (NCP) vs 22 (control), p-value = 1.4e-19] (**Figure 6a**). Since Phalloidin binds to and stabilizes F-actin filaments, the decrease in fluorescence intensity indicates F-actin depolymerization throughout the cell. Cytoskeletal reorganization has been linked to phosphatidylserine (PS) externalization (a process generally associated with apoptosis) suggesting a potential link between cellular paroxysm and plasma membrane integrity^35,36^.

**Figure 6:**
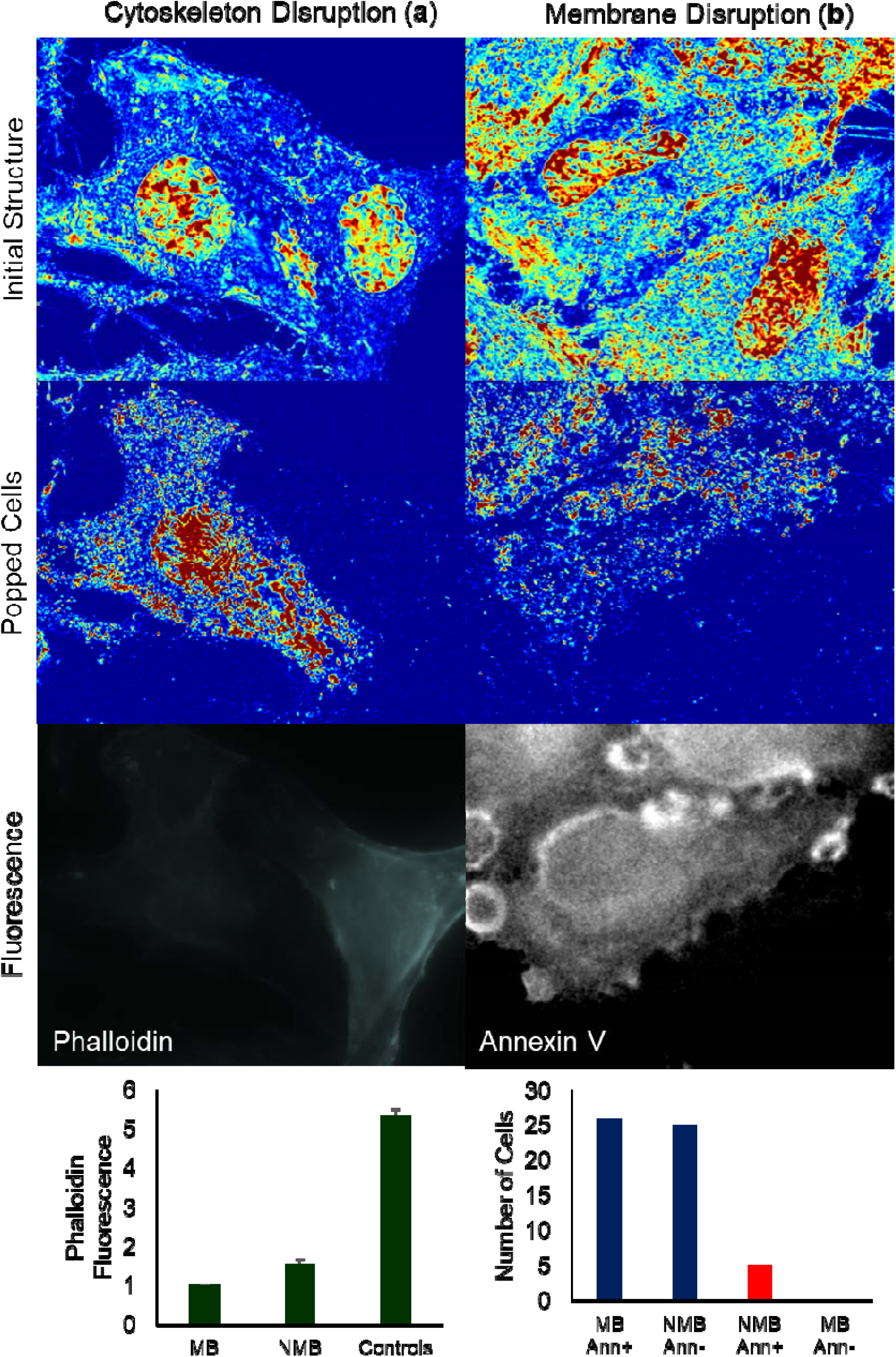
Cellular Paroxysm Biological Exploration. Exploration of cytoskeletal **(a)** and membrane **(b)** disruption during cellular paroxysm using Alexa Fluor 488 Phalloidin and Annexin V-FITC respectively. (row 1) Representative ∑_s_ maps of cells before UV irradiation. (row 2) Multiple m_f_ maps processed and combined to show which cells experienced the burst event throughout the experiment. (row 3) Representative fluorescence image of Fluor 488 Phalloidin **(a)** and Annexin V-FITC **(b)**. (row 4) Bar graphs showing the relationship between the burst event and the fluorescence measurements.

In healthy intact cells, PS is located on the interior of the plasma membrane. Annexin V was used to detect PS externalization since it cannot pass through an intact cell membrane, therefore, it will only bind to PS which has translocated to the exterior of the membrane (PS externalization). PS externalization is one of the earliest detectable processes in cells that have started to undergo apoptosis; although, there are some non-apoptotic causes of PS externalization such as engagement of immunoreceptors, ultrashort electric pulses, etc^36,37^. One of the most interesting biological results was a strong correlation between cellular paroxysm and Annexin V staining. 91% of the cells followed the trend where CP cells were Annexin V positive and NCP cells were Annexin V negative (26 cells CP and Annexin V positive, 25 cells NCP and Annexin V negative, 5 cells NCP and Annexin V positive, 0 cells CP and Annexin V negative) (**Figure 6b**). Finally, propidium iodide (PI) was used to detect poration of the plasma membrane. PI is an intercalating agent and fluorescent molecule that cannot enter cells with intact plasma membrane. All cells exhibited positive PI staining at the end of the experiment, indicating that UV caused poration of the cell membrane, but there is no correlation with cellular paroxysm (**Figure 7a)**. In summary, this burst of motion is connected to a disruption of the cytoskeletal and cell membrane structures, and may play a role in the early stages of apoptosis.

**Figure 7:**
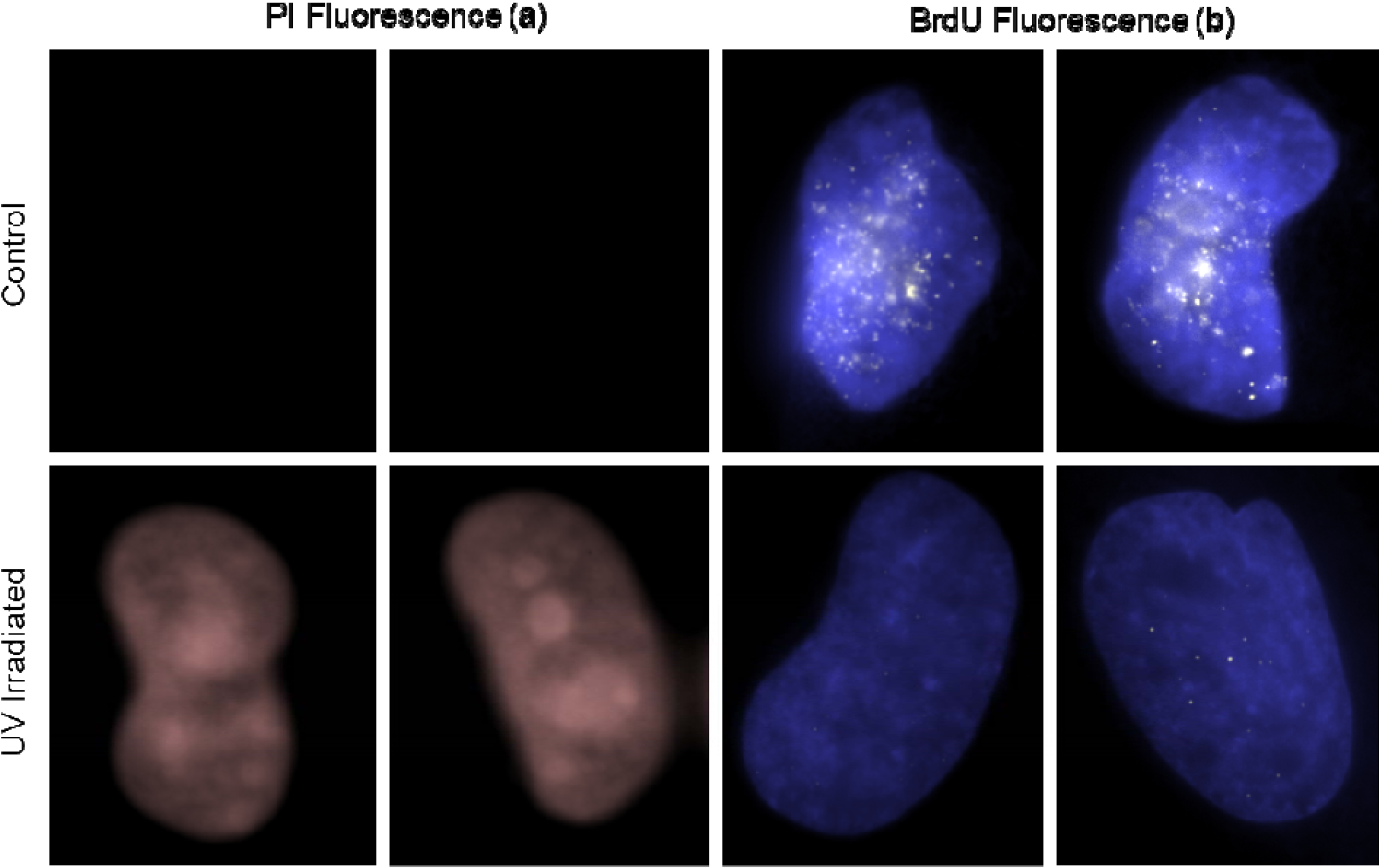
Membrane Poration and DNA Replication. **(a)** Representative fluorescence images of propidium iodide (PI) for control cells (top) and UV irradiated cells (bottom) show that pores form in the cell membrane in response to UV irradiation. **(b)** Representative fluorescence images of Bromodeoxyuridine (BrdU) for control cells (top) and UV irradiated cells (bottom) show that UV irradiation is stalling DNA replication.

UV irradiation is known to stall DNA replication. This halt of DNA replication seems to coincide with the arrest of nuclear motion detected with dual-PWS microscopy. To confirm these results, after UV irradiation (and arrest of nuclear motion), cells were stained with Bromodeoxyuridine (BrdU), a synthetic nucleoside that incorporates into freshly synthesized DNA in replicating cells. In control cells, we observed incorporation of BrdU into the nucleus, indicating active DNA replication (**Figure 7b**). In contrast, there was minimal incorporation of BrdU into the nuclei of UV irradiated cells. Additionally, we saw no difference in BrdU incorporation between CP and NCP cells. These results confirm that the UV irradiation in these experiments is stalling DNA replication, but this was independent of the cellular paroxysm occurring within the nucleus.

## Discussion

In this work, we present a novel imaging platform that combines the nanoscale structural measurements of PWS microscopy with a new technique for measuring nanoscale cellular dynamics. By capturing the temporal interference, this new technique quantifies cellular dynamics, such as the fractional moving mass and the diffusion coefficient. First, we validate this instrument comparing measurements of nanosphere phantoms to the analytical theory. As a biological control, we demonstrate that macromolecular motion ceases upon PFA fixation. We explored the structural and dynamic changes that occur in stem cells due to differentiation, demonstrating both increased variations in chromatin folding and molecular motion in the stem cell population. Finally, we studied the effects of UV irradiation and discovered a new biological phenomenon, where cells undergo a large rapid burst of motion prior to cell death.

Studying structural and dynamic alterations in stem cells in response to differentiation is an ideal model to explore the processes associated with cells that have undergone a major phenotypic transformation. Our lab has recently used live cell PWS to identify compounds that reduce variations in supra-nucleosomal chromatin folding and thus render cancer cells more susceptible to chemotherapy treatment^18^. The ability to find compounds that modulate the chromatin folding of cells along the stem continuum – that is, to find compounds that increase the rate of differentiation or that trigger de-differentiation – would prove to be crucial in better understanding the stem cell differentiation process and in guiding our search for compounds that could be used to treat patients with neurological disorders.

The results from the UV experiments provide for the first time high-temporal resolution, nanoscopic information about cellular motion during the UV irradiation. Interestingly, there are multiple novel phenomena observed that are the result of light mediated cellular stress. Initially, we see a homogenization of the chromatin structure, which is likely due to UV induced DNA fragmentation that is paired with an arrest in cytoplasmic and nuclear motion. The observed arrest in motion in the cytoplasm converges with actin depolymerization (**Figure 6a**) (consistent with decreased cytoplasmic Σ*_s_*) and poration of the cell membrane (**Figure 7a**). After cells reach a near static state, a stochastic eruption of motion (cellular paroxysm) forms across the entire cell. While the cellular paroxysm is asynchronous from cell to cell across the field of view, it frequently synchronizes for cells in contact with each other. Temporal analysis of events within individual cells show that these events can occur across the entire cell as far as 30 μm apart in under 35ms.

To be synchronous and diffusive in nature, a molecular regulator would need to move at an incredibly high speed 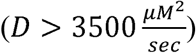 to facilitate this process. Molecularly, the cellular paroxysm correlates with PS externalization and actin depolymerization. The correlation with actin depolymerization has two possible explanations. The first explanation is that part of this event is an additional rapid depolymeriation of actin. As actin accounts for 5–10% of all protein within the cell, spontaneous alteration of that entire structure would produce a significant movement of mass throughout the entire body of the cell^38^. Alternatively, actin depolymerization is a precursor to this event, and therefore cells which depolymerize actin faster will undergo the cellular paroxysm first. Interestingly, the cellular paroxysm also takes place within the nucleus indicating either a direct integration of chromatin motion with actin depolymerization at short timescales or a separate determining mechanism.

In addition to the coupling of cellular paroxysm with actin depolymerization, there is a strong correlation with PS externalization (100% of CP cells are Annexin V positive and 83% of NCP cells are Annexin V negative). This suggests a synchronous coupling between cellular paroxysm, cytoskeletal organization, and membrane structure at short timescales. Further, as PS externalization is highly correlated with apoptosis, this suggests that cellular paroxysm could be a never before seen early event in UV mediated cell death. With respect to the cells that were uncorrelated (9% of cells), these cells displayed positive Annexin V staining without an accompanying cellular paroxysm. Notably, there were no CP cells that displayed negative Annexin V staining. The likely explanation is that these uncorrelated cells did actually experience cellular paroxysm, but the events were not captured. One reason that we may have missed a few of these events is that there is a short time gap (<20 seconds) between the completion of our final measurement in these experiments and the UV light being turned off. It is difficult to speculate on the magnitude of fractional moving mass that a PS externalization event would cause as the exact physical mechanism is unknown, but the independent motion of PS would produce a weaker signal than largescale actin depolymerization. These experiments suggest that both of these molecular transformations are linked to a structural transformation in the cell at short timescales.

## Conclusion

In summary, we have developed a system that enables label-free live cell measurements of nanoscale structure and macromolecular motion with millisecond temporal capabilities, derived new theory for measurements of temporal interference, and validated the system using experimental phantoms. Then the functionality of dual-PWS was explored through measurements of cellular fixation, stem cell differentiation, and the cellular response to UV irradiation. Through the stem cell experiments, we observed alterations in chromatin ultrastructure, as well as the fractional moving mass and diffusion of chromatin in response to differentiation. We found that UV-irradiation induces homogenization of chromatin structure, while decreasing the fractional moving mass and chromatin diffusion. The exploration of UV-irradiation led to the discovery of a new phenomenon, a near-instantaneous burst of motion (cellular paroxysm) that may be an early step in the process of UV induced cell death due to its correlation with PS externalization.

To our knowledge, this is the first time this phenomenon has been observed and it exemplifies some of the unique features and advantages of the dual-PWS microscope, making it uniquely suited to analyze macromolecular processes such as this one. As a high-speed imaging technique, dual-PWS can track individual cells in real time, enabling the identification of asynchronous changes (like the cellular paroxysm), comparisons of the same cells, before, after, and during treatments, and detection of alterations in sub-cell populations. By utilizing the interference phenomenon, dual-PWS images without labels, avoiding the artifacts and cytotoxicity associated with dyes. It is important to note that PWS is high throughput, and thus it can measure dozens of cells in just seconds. These key features of dual-PWS enable new exploration of critical biological questions about macromolecular behavior in live cells.

This technique could provide valuable information about structure-function relationships within the nucleus, especially for the important processes of repair, replication, and transcription. In particular, it can facilitate investigation about the spatio-temporal biological regulators of chromatin, such as: does the cytoskeleton or the nuclear lamina regulate chromatin, and what role does the intranuclear electrostatic environment play? With respect to human diseases, this technique allows investigation into therapeutics that regulate chromatin structure and dynamics in fields such as regenerative medicine, cancer, and infectious diseases. While chromatin is of particular interest as the regulator of transcription, replication, and repair, this technology can be used to study any biological systems, such as cytoskeletal dynamics during the processes of cell division, muscle contraction, or carcinogenesis.

## Methods

### Microscope Design and Data Acquisition

The dual-PWS system consists of a commercial microscope base (Leica DMIRB) with a white light LED source (X-Cite 120LED), 63x oil immersion objective (Leica HCX PL APO, NA 1.4 or 0.6), spectral filter (CRi VariSpec LCTF), and CCD camera (Hamamatsu Image-EM CCD). Light from the source is focused onto the sample with an illumination NA of 0.55, the backscattered light is collected by passing it through the spectral filter and imaged at the CCD. Multiple wide-field monochromatic images are obtained for each acquisition and stored in a three-dimensional image cube as described below.

Structural measurements are collected by imaging the backscattered light across a range of wavelengths (500–700nm) to produce a spectral data cube, I(λ, x, y), where λ is the wavelength and (x, y) correspond to pixel positions. Each frame within a spectral data cube is acquired with a 35ms exposure, and the total cube can be obtained in under two seconds depending on the number of wavelengths collected^39^. Similarly, dynamics measurements are obtained by imaging the backscattered light at a single wavelength (550nm) over a period of time, to produce a temporal data cube, I(t, x, y), where t is time and (x, y) correspond to pixel positions. To isolate the dynamics originating from our biological sample as opposed to experimental noise (i.e. temporal lamp fluctuations), 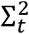 is measured from an empty sample and subtracted from our biological sample’s 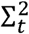. For this study, temporal measurements consisted of at least 201 frames acquired with 1.2ms, 32ms, or 35ms exposures each for a total time of ranging between 240ms to 7 seconds. In theory, the rate of acquisition and the number of frames acquired can be adjusted to match the biological phenomenon of interest as explained below. In the current configuration, the temporal and structural measurements are performed sequentially in <15 seconds.

### General Diffusion Analysis

Diffusion coefficients were calculated in the following manner:

1) Temporal interference data cube is loaded and normalized for lamp intensity, exposure time, and dark counts.
2) The temporal mean is subtracted from each pixel so that the temporal oscillations fluctuate around zero.
3) The temporal autocorrelation function (ACF) is calculated at each pixel.
4) ACFs with low SNR are removed. This is determined by pixels where the first point of the ACF is less than 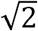 multiplied by the first point of the ACF of the background (region without cells).
5) The average ACF of a background sample (no cell or spheres) is subtracted from each individual ACF.
6) The ACF at each pixel is normalized so that the first point equals one.
7) Negative values are removed in order to calculate the natural log of the ACF at each pixel.
8) The natural log of the ACF is averaged across all pixels. The mean ACF can be obtained by inversing the natural log using the exponential function, although this is not necessary to calculated *D*.

a. In order to generate diffusion maps, *D* must be calculated at each pixel so the natural log of the ACF cannot be averaged. To account for the increased noise, the ACF is filtered spatial with a Gaussian filter and then temporally with a moving average filter. The SNR threshold was also increased to 2* background signal.
9) The slope of the natural log of the ACF is calculated from the first two points to acquire the decay coefficient of the mean ACF.
10) D is calculated from the decay coefficient using the following formula. 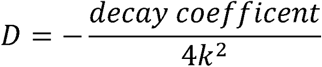

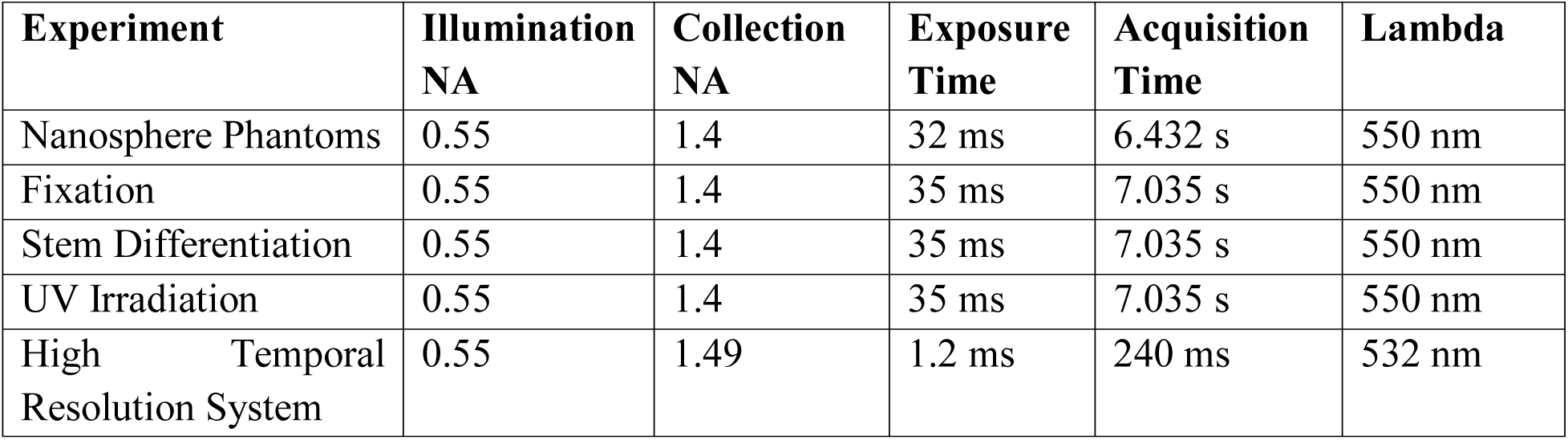
Experimental Imaging Parameters

### Statistical Analysis

All t-tests are two tailed, and heteroscedastic unless otherwise noted as paired. All error bars are standard error of the mean (SEM). All n and p-values are reported in the text near the relevant comparison. All the parameters were calculated using Microsoft Excel (Microsoft Corporation, Redmond, WA).

### Cell Culture

All experiments (except for stem cell experiments) were performed with HeLa cells. HeLa cells (ATCC, Manassas Virginia) were grown in Gibco® formulated RPMI-1640 Media (Life Technologies, Carlsbad California) supplemented with 10% FBS (Sigma Aldrich, St. Louis Missouri) and grown at 37°C and 5% CO_2_. All of the cells in this study were maintained between passage 5 and 20. Cell were grown in 50mm petri dishes with uncoated size 0 or 1 glass coverslip bottoms (MatTek, Ashland Massachusetts). Petri dishes were seeded with between 10,000 and 50,000 cells in 2ml of the cell appropriate media at the time of passage. Cells were allowed at least 24 hours to re-adhere and recover from trypsin-induced detachment. Imaging was performed when the surface confluence of the slide was between 40–70%.

### Stem Cell Culture and Differentiation

Human mesenchymal stem cells (hMSCs, ATCC) were cultured in Dulbecco’s Modified Eagle Medium (DMEM), with 4.5 g/L glucose, and supplemented with 10% FBS and 5ml 10x penicillin-streptomycin. For differentiation studies, cells were plated 1.5×10^4^ cell/ml in 24 well glass-bottom PWS plates. After 2 days, regular DMEM was switched to hMSC Osteogenic Differentiation Medium (Lonza) containing B-glycerophosphate, ascorbate and dexamethasone. Media was changed every other day and cells were imaged on day 4 post-induction. To confirm differentiation, ALP colorimetric assay was performed on fixed hMSCs after imaging live with dual-PWS.

### Alexa Fluor 488 Phalloidin Experiment

Cells were irradiated with UV for ~12 minutes. First, cells were washed with PBS. Then, cells were fixed with 3.6% paraformaldehyde solution for ~10 minutes. A permeabilization/blocking step was performed with 0.1% Triton X-100, 1% bovine serum albumin in PBS for 5 minutes. Finally, cells were stained for 20 minutes with 40x dilution Alexa Fluor 488 Phalloidin (Thermo Fisher Scientific).

### Annexin V

Cells were irradiated with UV for ~12 minutes. Then, 10% of the media was removed and replaced with an equivalent volume of 10X Annexin V Binding Buffer. Next, cells were stained with ~100x dilution of Annexin V-FITC Conjugate for 10 minutes. For some runs, staining was performed before the UV irradiation, so that multiple runs could be accomplished within the same dish. Annexin V Binding Buffer and Annexin V-FITC Conjugate come from Annexin V-FITC Early Apoptosis Detection Kit from Cell Signaling Technologies.

### Propidium iodide

Cells were irradiated with UV for ~12 minutes. Then, 10% of the media was removed and replaced with an equivalent volume of 10X Annexin V Binding Buffer. Next, cells were stained with ~9x Propidium Iodide for 10 minutes. For some runs, staining was performed before the UV irradiation. Annexin V Binding Buffer and Propidium Iodide came from Annexin V-FITC Early Apoptosis Detection Kit from Cell Signaling Technologies.

### BrdU

Cells were irradiated with UV for ~15 minutes. Then, BrdU was added to a concentration of 0.03 mg/mL and incubated at 37C for 30 minutes. The cells were then fixed and stained with a secondary antibody according to the manufacturer’s protocol found at https://www.cellsignal.com/contents/resources-protocols/immunofluorescence-protocol-for-labeling-with-brdu-antibody/5292-if

## Acknowledgements

This work was supported by fellowships and grants from the National Institutes of Health grant R01-GM105847, R01 CA200064, R33CA225323, 1R01CA228272, R01CA225002, R01EB016983, R01CA165309 and the National Science Foundation grant CBET-1240416.

## Contributions

S.G. designed the study, performed experiments, analyzed data, developed the theory, validated the system, and wrote the manuscript. L.M.A. designed the study, performed experiments, analyzed data, and wrote the manuscript. L.C. developed the theory, and reviewed the manuscript. A.E. performed experiments, and reviewed the manuscript. A.E., D.Z., W.W. validated the system, and reviewed the manuscript. G.M.B., S.M. performed experiments, analyzed data, and reviewed the manuscript. J.E.C., A.D.S., H.S., J.F.M., G.A.A., I.S. designed the study, and reviewed the manuscript. V.B. designed the study, developed the theory, and reviewed the manuscript.

## Competing Interests

There are no competing interests to report.

## Video Legends

**Video 1** Ultraviolet Irradiation Video. Representative m_f_ (top) and ∑_s_ (bottom) videos for cells undergoing UV (right) and non-UV (left) irradiation.

**Video 2** Motion Burst Timing Video: Video showing the timing of the motion burst event for each pixel within a representative cell. The spatial distribution of this timing shows that the initial motion occurs simultaneously within a single 35ms frame on opposite ends of the cell.

